# Identifying Naturalistic Movies from Human Brain Activity with High-Density Diffuse Optical Tomography

**DOI:** 10.1101/2023.11.27.566650

**Authors:** Zachary E. Markow, Kalyan Tripathy, Alexandra M. Svoboda, Mariel L. Schroeder, Sean M. Rafferty, Edward J. Richter, Adam T. Eggebrecht, Mark A. Anastasio, Mark A. Chevillet, Emily M. Mugler, Stephanie N. Naufel, Allen Yin, Jason W. Trobaugh, Joseph P. Culver

## Abstract

Modern neuroimaging modalities, particularly functional MRI (fMRI), can decode detailed human experiences. Thousands of viewed images can be identified or classified, and sentences can be reconstructed. Decoding paradigms often leverage encoding models that reduce the stimulus space into a smaller yet generalizable feature set. However, the neuroimaging devices used for detailed decoding are non-portable, like fMRI, or invasive, like electrocorticography, excluding application in naturalistic use. Wearable, non-invasive, but lower-resolution devices such as electroencephalography and functional near-infrared spectroscopy (fNIRS) have been limited to decoding between stimuli used during training. Herein we develop and evaluate model-based decoding with high-density diffuse optical tomography (HD-DOT), a higher-resolution expansion of fNIRS with demonstrated promise as a surrogate for fMRI. Using a motion energy model of visual content, we decoded the identities of novel movie clips outside the training set with accuracy far above chance for single-trial decoding. Decoding was robust to modulations of testing time window, different training and test imaging sessions, hemodynamic contrast, and optode array density. Our results suggest that HD-DOT can translate detailed decoding into naturalistic use.

## 1. INTRODUCTION

Modern neuroimaging modalities, particularly functional MRI (fMRI), can decode remarkably detailed information about novel stimuli and actions experienced or imagined by human subjects. The identities of thousands of images can be decoded along with object classification, and more-ambitious paradigms can reconstruct visual scenes or sentences.^1–8^ In many cases, decoding paradigms leverage encoding models that reduce the full, all-possible, stimulus space down into a reduced yet generalizable set of stimulus features. While the complexity of the tasks continues to increase at a remarkable pace, these devices are either non-portable, like fMRI, or invasive, like electrocorticography (ECoG), limiting their applicability in chronic brain-computer interfaces or naturalistic settings. Wearable, non-invasive, but lower-resolution and lower-signal-to-noise devices such as electroencephalography and functional near-infrared spectroscopy (fNIRS) have generally been limited to only template-based, non-generalizable decoding with very little detail.^9–12^ Herein we develop and evaluate model-based decoding with high-density diffuse optical tomography (HD-DOT), a higher-resolution expansion of fNIRS that has shown promise as a surrogate for fMRI.^13–17^ We decode the identities of novel, previously unseen movie clips among 4-40 options with a motion-energy encoding model,^2^ expanding on previous optical studies that used template-based decoding or simpler methods and thus were not able to decode outside of the training stimuli.^9–12,18^

Neural decoding infers partial information about a stimulus, state, or imagined action from measurements of brain activity evoked by that stimulus, state, or action.^8^ This decoding enables key scientific and clinical applications, such as testing whether particular brain regions carry information about specific stimulus features and forming brain-computer interfaces that enable people to perform augmented communication or control a device with thought instead of explicit movement. Over the last decade, fMRI, ECoG,^6^ and intracortical recordings^7^ have achieved remarkable and complex stimulus decoding capabilities. For example, previous fMRI studies identified which of 120 images a subject viewed,^1^ reconstructed estimates of movies viewed,^2,4^ classified objects seen,^3^ and semantically reconstructed sentences heard or covertly spoken by subjects.^5^

Fundamental to the more ambitious decoding paradigms, that decode between previously unseen visual or auditory stimuli, or reconstruction of stimuli, is the creation (through training) of parametric models that predict brain activity from stimulus features or vice versa.^1–3,5^ Parametric decoding allows decoding of stimuli outside the training set, by addressing the combinatorial explosion of possible stimuli, even at the scene, word, and sentence level. Since an all-encompassing training set of all possible stimuli is impossible to acquire, parametric approaches work to compress the possible stimuli into reduced feature spaces, and then learn the mapping between features and brain imaging measures.^8^

Although fMRI, ECoG, and intracortical recordings have achieved remarkable, complex stimulus decoding capabilities, the logistics of these modalities make them impractical as chronic brain-computer interfaces. For fMRI, the subject would need to remain in a bore that is exceptionally non-portable and expensive. For ECoG and intracortical recordings, the technology is invasive requiring at a minimum removal of a portion of the skull. In contrast, electroencephalography (EEG) and sparse functional near-infrared spectroscopy (fNIRS) are lighter-weight, non-invasive portable tools. However, sparse channel-count systems like EEG and conventional fNIRS with <100 channels, have demonstrated far more-limited decoding capabilities than fMRI and ECoG.^9–12^ High-density diffuse optical tomography (HD-DOT) systems, with ∼128 sources and detectors that combine for >2000 usable source-detector pairs (channels), has the potential to address the challenges of translating model-based decoding out of the fMRI scanner and into naturalistic environments. HD-DOT systems are non-invasive and have spatial resolution approaching group-averaged fMRI.^13–16,19–21^ While most fiber-based HD-DOT systems are not wearable and only modestly portable, the technology is currently going through transformation towards full-head wearable systems.^19–21^ Because decoding performance largely scales with information content, the improved spatial resolution of HD-DOT suggests that it will enable neural decoding capabilities closer to fMRI than sparse fNIRS.

Although modest HD-DOT decoding studies have been conducted in the past, these studies were limited to decoding the same stimuli used in training and they have yet to attempt more general out-of-sample decoding, where novel stimuli not used in training are decoded, on a single-trial basis.^18,22^ Here, we evaluate the feasibility model-based visual decoding with HD-DOT. While we use a fiber-based HD-DOT system, we anticipate that these results will translate to future wearable HD-DOT systems, as long as the systems match the critical performance metrics of fiber-based HD-DOT.^21^ Visual decoding was one of the first steps in the chain of advances in fMRI decoding during the last decade,^1,2^ and here we begin translating that progress to HD-DOT.

## 2. RESULTS

### 2.1. Validating Data Quality and Response Repeatability

In all decoding tasks, the goal was to identify which movie clip the subject was viewing at a given time, using diffuse optical tomography (DOT) data. To perform stimulus decoding, we employed a UHD-DOT system to image brain activity in occipital cortex in healthy adults while they watched four 90-second, audio-free movie clips 4-10 times per clip, in each of 4 imaging sessions (**Fig. 1A-B**).

**Figure 1.**
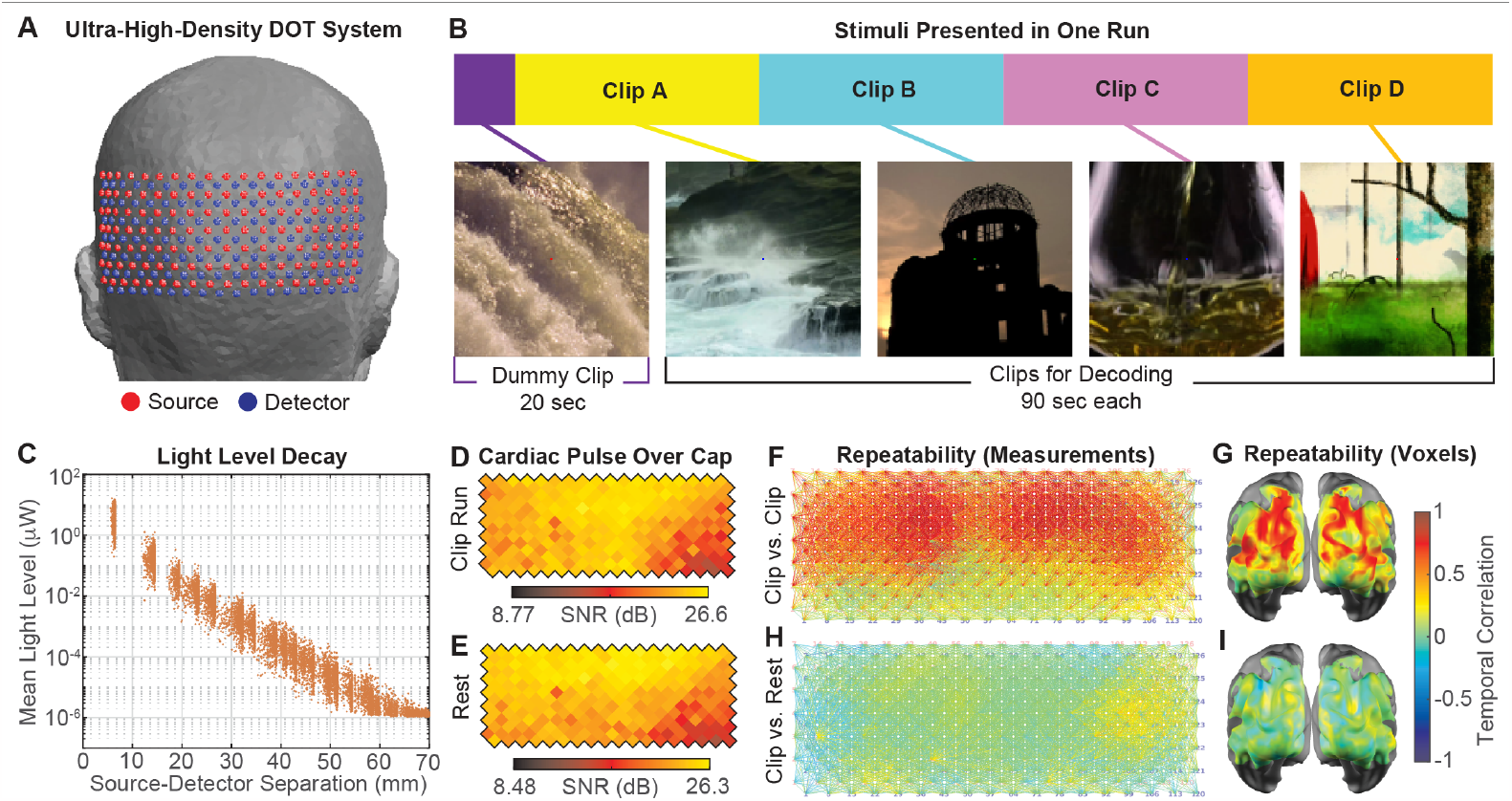
UHD-DOT data quality during movie clip viewing. (**A**) UHD-DOT optode grid and placement on head. (**B**) Stimulus presentation in one example movie run. (**C**) Light level decays log-linearly with source-detector distance stretching over 7 orders of magnitude. (**D-E**) Ratio of cardiac pulse strength to background signal strength in channels with 21-35-mm source-detector distance shows excellent hemodynamic signal fidelity across the cap. (**F-G**) Correlation between the brain’s average imaged response to one clip from first half vs. second half of the runs within a session, for each channel (**F**) and voxel (**G**). (**H-I**) Correlation between the same average response vs. similarly averaged brain activity during resting-state (fixation cross) epochs. The far-higher correlations in panels **F-G** than **H-I** indicate that imaged clip responses were repeatable compared to stimulus-unrelated variance and correlation. Panels **C-I** are exemplary data from one imaging session in one subject.

Decoding performance relies on raw data quality, signal-to-noise ratio, and repeatability of clip-evoked brain responses. Therefore, following data collection on 12 subjects, we evaluate raw data quality, including checking that the light levels in each measurement channel decayed log-linearly with source-detector separation and computed the cardiac pulse signal strength above background physiology before filtering (**Fig. 1C-E**).^22^

To evaluate the repeatability and similarity of the brain’s responses to each clip, we computed the correlation between the average brain response to each clip from the first and second half of each imaging session (**Supplementary Figs. S1, S2**). In the 6 subjects with acceptable raw data quality and clip response repeatability (**Supplementary Fig. S3**), we found the correlations between the average responses to the same clip (matched) where much higher than correlations to a gray screen (rest), indicating that these clip responses were repeatable (**Fig. 1F-I**), suggesting these data sets were good candidates for evaluating model-based decoding.

### 2.2. Parametric Encoding Model Predicts Brain Responses

Parametric model-based decoding outside the training set requires an encoding model that maps a common set of features in stimuli to brain responses at each voxel. Therefore, we used the brain responses to two clips within a session (training clips) to fit a parametric encoding model that predicts the brain’s response to an arbitrary clip as a weighted sum of wavelet motion energy features (**Figs. 2, 3**; **Supplementary Fig. S4**).^2,23–25^ Our encoding model is 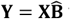 where **Y** (time-by-voxels) is the measured brain response, **X** (time-by-features) is the extracted feature representation of a movie sequence convolved with a hemodynamic response function, and 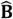 (features-by-voxels) is the mapping (weights) between features and voxels. The goal of model training is to estimate 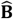, which we do by using ridge regression to calculate the inverse of **X**_train_ and then calculate 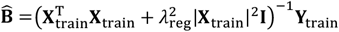. The regression regularization parameter *λ*_reg_ was scaled by the squared norm of **X**_train_ and set by cross-validation. From this model, the predicted brain response **Y**_pred_ to any clip can be computed by similarly decomposing the clip into its feature time-traces **X**_test_ and multiplying by the weight matrix 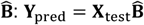.

**Figure 2.**
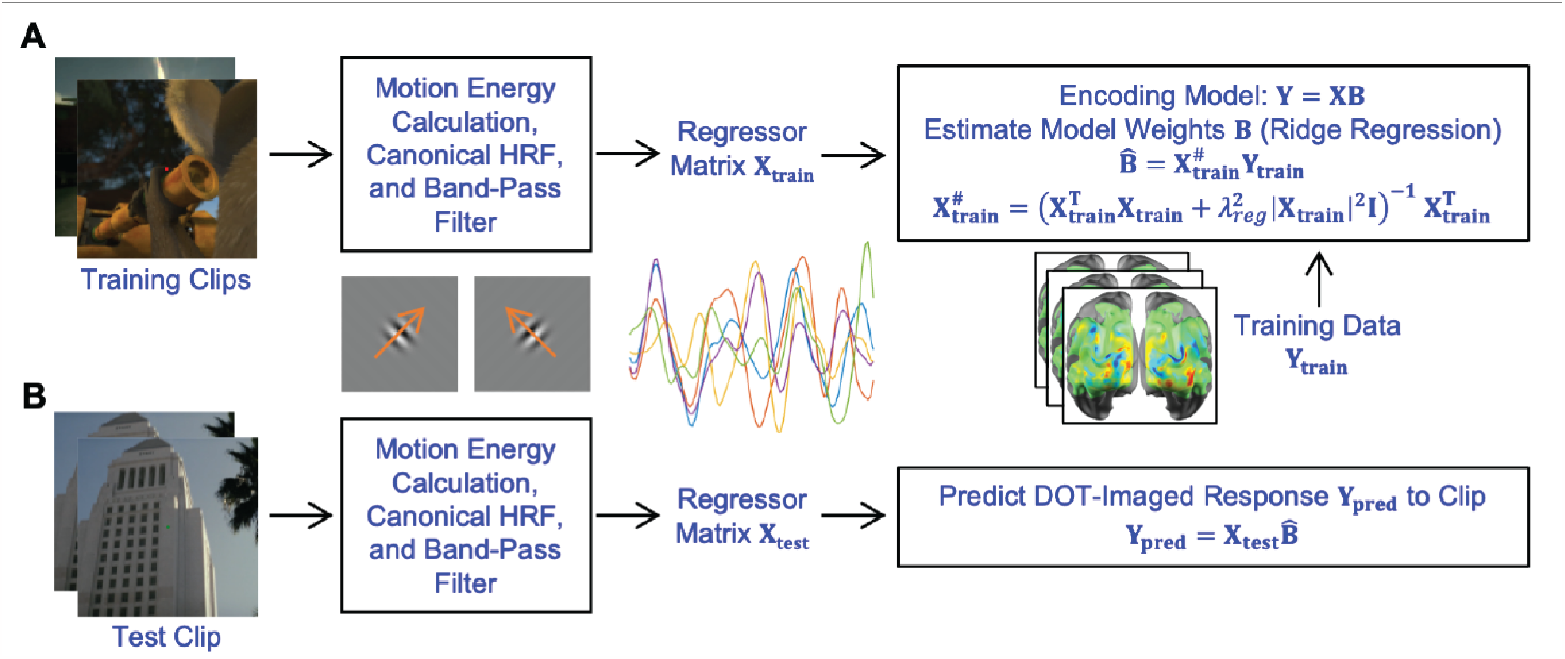
Building the parametric encoding model. (A) Encoding model treats the image time trace in each voxel as a weighted sum of feature time traces in the movie. Here, each feature measures the strength of motion at a given speed, orientation, location, and spatial frequency in the movie. Two clips’ trials are used to train (fit) the encoding model, while the other clips’ trials are held out to test prediction and decoding accuracy. For training, the training clips are decomposed into their wavelet motion energy feature time traces. Then these traces are convolved with a canonical hemodynamic response function (HRF) and band-pass filtered as in DOT data preprocessing to yield the feature regressors. Ridge regression then estimates the model weights 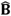 from the imaging data and feature regressors. (B) After training, the same pipeline and encoding model can calculate the predicted brain response to any arbitrary clip.

**Figure 3.**
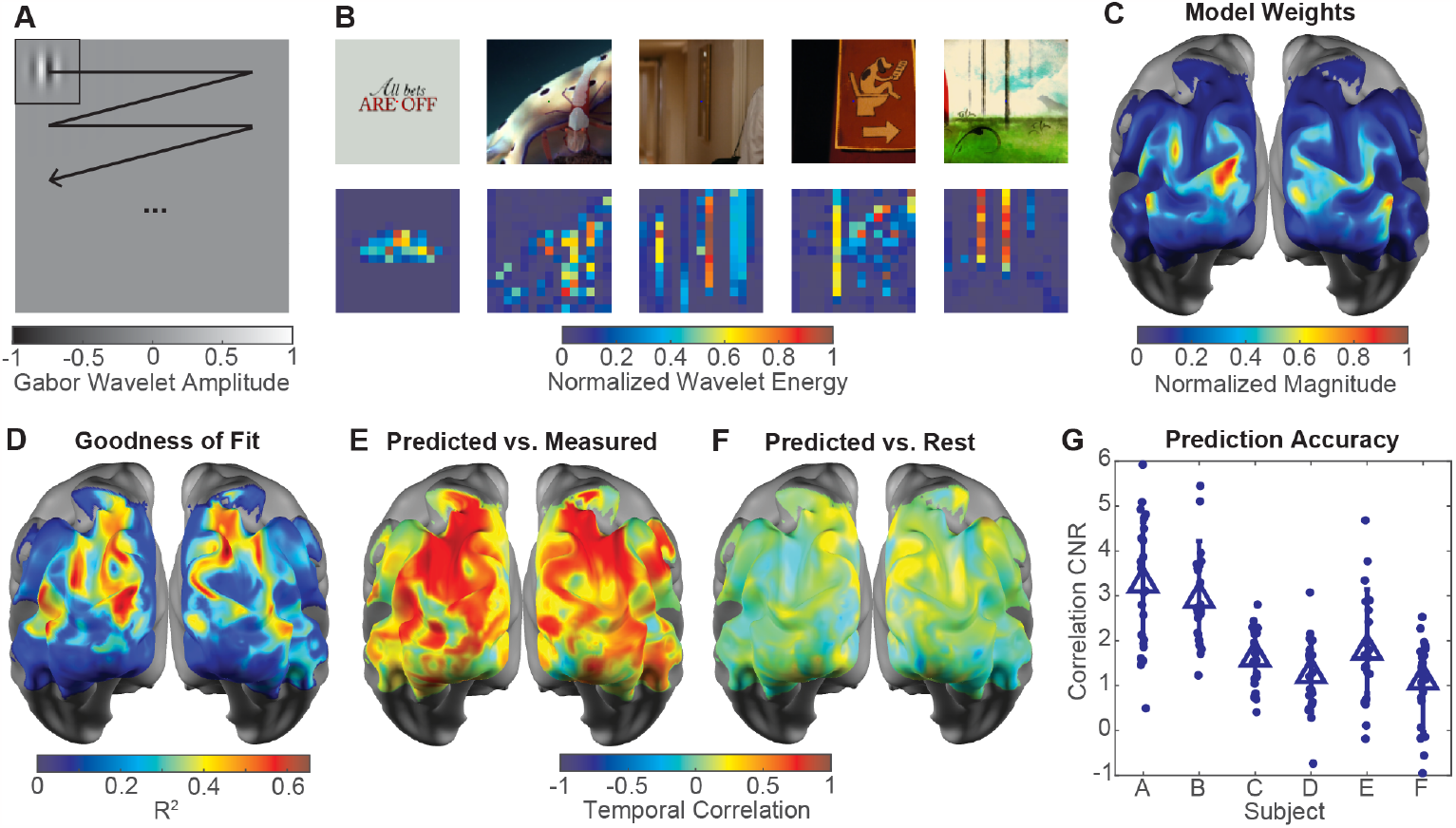
Encoding model validation and analysis. (A) Example wavelet motion feature. (B) Example feature’s strength at 16×16 different locations in different movie frames. (C) RMS magnitude of encoding model feature weights at each brain location indicates larger sensitivity in occipital regions that respond strongly to visual stimuli and fall within areas where the imaging system has high SNR and sensitivity. (D) Encoding model explains over 30% of the variance in the imaged brain responses in many voxels by the *R*^2^ metric. (E-F) Correlation over time between predicted response and actual data for held-out test clip exceeds 0.5 and far-exceeds correlation between predicted response and resting-state data in many voxels, indicating high prediction accuracy. (G) Distribution of these prediction-vs-clip-data correlations over space and time far-exceeds the distribution of prediction-vs-rest-data correlations for most sessions and train-test splits, as measured by Cohen’s d statistic (CNR) and bootstrap test.

To check that the model predicted brain responses reasonably accurately, we computed the correlations between the predicted and actual brain responses for each non-training clip as well as the correlations between the predicted response to one clip and actual brain responses to resting-state data or a different clip. We found that the model predicted clip responses far more-accurately than the “null” responses from the resting-state data and mismatched clips (**Figs. 3E-G, 4A-B; Supplementary Fig. S5**), with *p*<10^−4^ and effect size 1.31 for predicting clip responses vs. rest responses.

**Figure 4.**
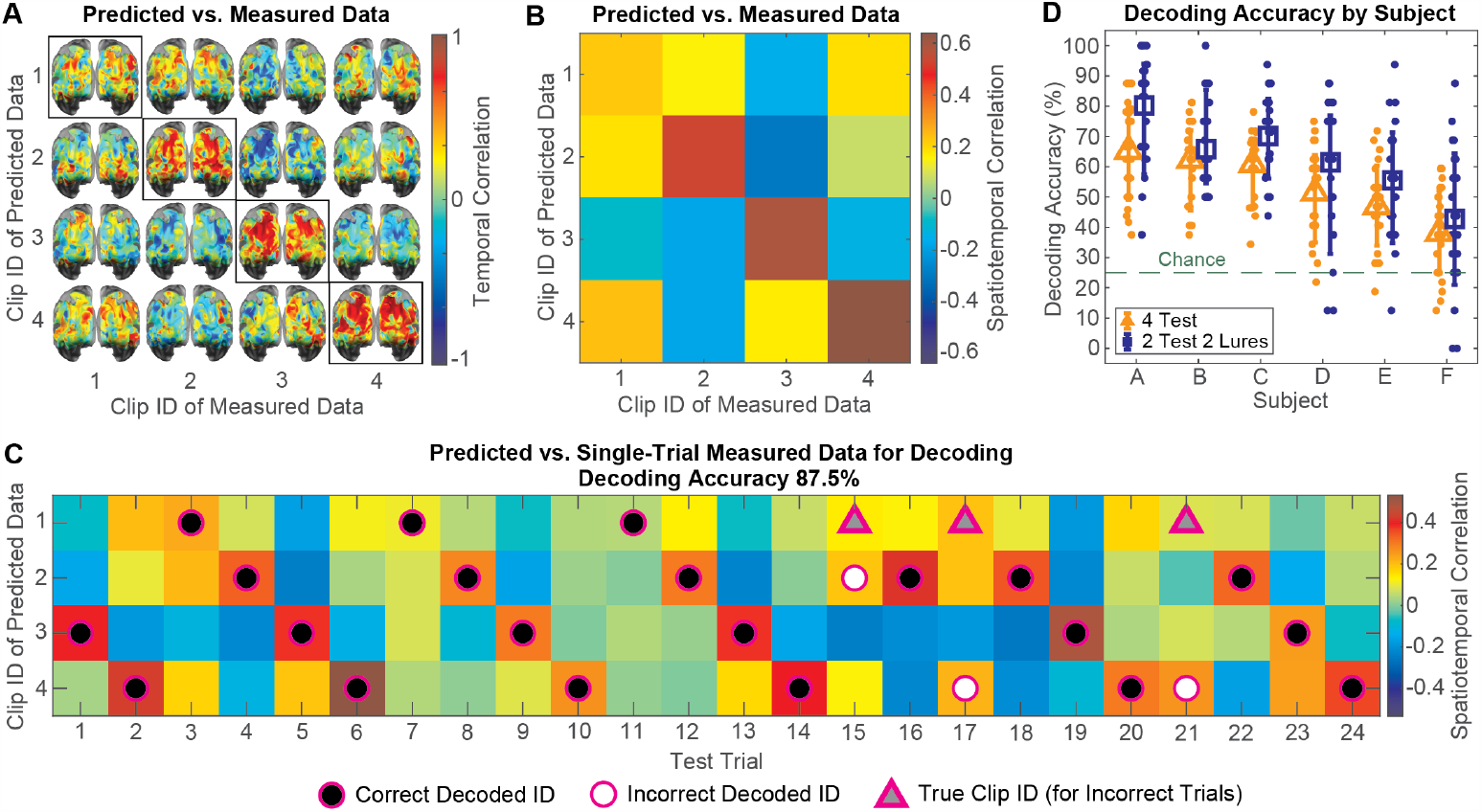
Model-based decoding. (A) Correlation over time between predicted response and actual data for the same/”matched” test clip (on-main-diagonal maps) or for a different/”mismatched” test clip (off-diagonal maps) for one session. (B) Similar correlations, but over both space and time. Greater correlations along the main diagonal than most off-diagonal maps/elements indicate accurate, discriminable predictions by the encoding model among clips. (C) Correlation between each test clip’s predicted response and the imaging data in each trial enables maximum-correlation decoding of clip identity shown in each trial. (D) Decoding accuracy in each session for 6 train-test splits of the clips, grouped by subject. Large markers and error bars indicate mean ± lower and upper standard deviation over sessions and train-test splits. Accuracies are shown for decoding among 4 half-clips (orange triangles) and among 2 original + 2 lure clips (blue squares).

### 2.3. Model-Based Decoding of New Clip Identities

Decoding starts with new data from clips previously held out from the encoder. The new responses are then compared to encoding model predictions to infer which clip was shown. To decode clips that were not in the training set, we programmed a decoder to calculate the correlation between the data and hypothetical clips and then pick the clip with the highest correlation (**Fig. 4C, Supplementary Fig. S6A**).

To evaluate decoding among 4 clip options we split each of our original 90-second clips in half, which gave 8 shorter clips total, 4 of which were test clips (**Supplementary Fig. S1**). Under these conditions, the maximum-correlation decoder achieved 54±16% accuracy (average ± standard deviation) over the 6 subjects with adequate raw data quality and repeatable clip responses. Those subjects’ 4 sessions, and 6 ways of separating the clips into training and test sets (“train-test splits”) had 25% chance level, thus our average decoding performance was well above chance (**Fig. 4D**, orange triangles, *p*<10^−5^, effect size 1.81). To enable training and evaluation on longer clip segments up to 75 seconds, we also performed a modified 4-clip decoding task in which the decoder was trained on 2 of the original 4 (non-split) clips, but the decoder guessed among the other 2 original clips plus 2 additional “lure” clips. These “lure” clips were never presented to subjects, but their predicted responses were generated with the encoding model (**Fig. 4D**, blue squares). These conditions yielded 62±22% average decoding accuracy, which far-exceeded the 25% chance level (*p*<10^−5^, effect size 1.68).

### 2.4. Increasing Task Difficulty: Decoding Between Sessions, with Less Data, and Among More Clips

Encouraged by these results, we next evaluated decoding under several conditions that arise in decoding applications but make accurate decoding more challenging. These manipulations also provide a negative control for the decoding results because the manipulations should degrade the accuracy. To evaluate feasibility of not retraining the decoder every day, we performed “between-session” decoding, in which we formed the decoder’s feature weights 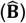 and predicted responses from one session and used these to decode the clip identities in another session, which gave 61±23% accuracy, statistically identical performance to within-session accuracy (**Fig. 5A**, *p*=0.087). Next, to assess how quickly decoding accuracy degrades with shorter durations of data, we used progressively smaller time windows of each clip response to compute the correlations for the decoder, finding that decoding accuracy declined very little from 75-35 sec and declined but remained statistically above chance from 35-10 sec (**Fig. 5B**). To evaluate feasibility of decoding among many more clips, we gradually increased the number of “lure” clips whose predicted responses the decoder was allowed to choose from, yielding accuracies that remained far above chance as the number of lure clips increased (**Fig. 5C**).

**Figure 5.**
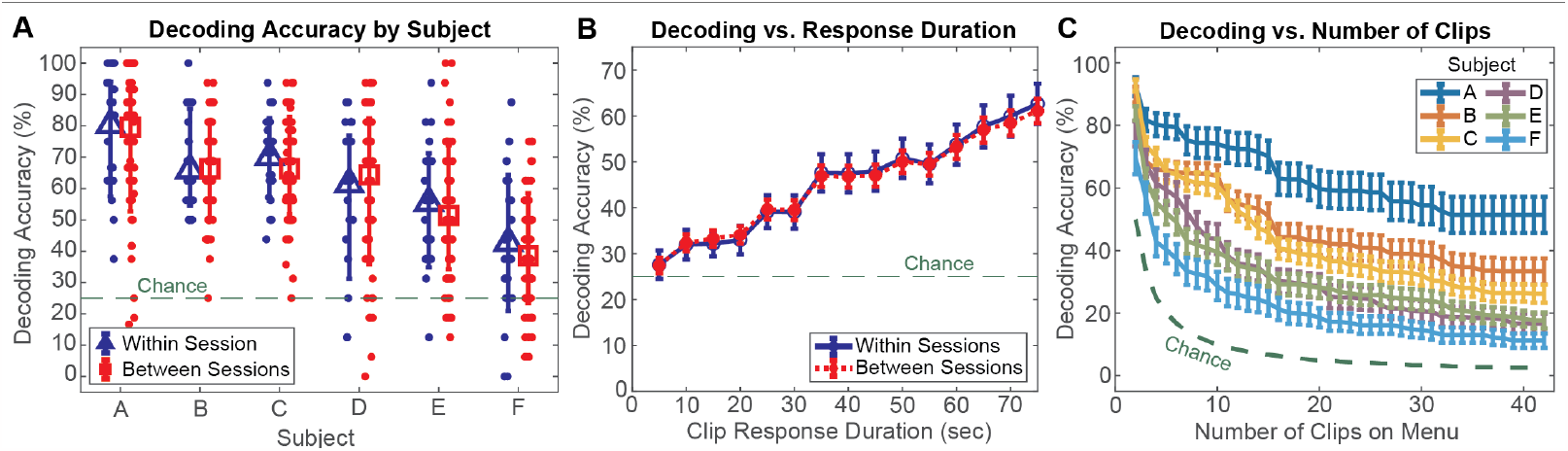
Decoder robustness. Paradigm manipulations indicate decoder robustness and generalizability across sessions, data duration, and number of clips. For within-session decoding, training and test trials were taken from the same imaging session, whereas for between-session decoding, training trials were taken from the same subject but a different session from the test trials. (A) Within-session and between-session decoding accuracy over sessions (for within-session) or session pairs (for between-session) and 6 train-test splits. Large markers and error bars denote mean ± lower and upper standard deviation. (B) Decoding accuracy declines as expected but remains above chance as the time window used for decoding shortens. Markers and error bars denote mean ± standard error over subjects, sessions, and 6 train-test splits. (C) Decoding accuracy declines but remains above chance, far above chance for some subjects, as the number of lure clips increases. Error bars denote standard error over sessions and train-test splits.

### 2.5. Decoding with Sparser Imaging Grid and Different Hemoglobin Contrasts

To assess the possibility of using sparser HD-DOT arrays, we reconstructed images of brain activity from an ordinary high-density subset of the UHD-DOT array, and we repeated decoding on those images, obtaining lower average accuracy than the UHD-DOT array (**Fig. 6A-B**): 62±22% accuracy for UHD vs. 56±15% accuracy for HD-subset (*p*<10^−4^, effect size 0.31). This finding was consistent with previous work indicating superior image quality and higher template-based decoding performance for UHD-DOT than its HD-subset.^22^ We also performed a simpler, template-based decoding task, in which we designated trials in the first and second half of each session as training and test trials, respectively, block-averaged the training trials to form “templates”, and used a maximum-correlation classifier to guess which of the 4 clips was viewed in each test trial (**Supplementary Figs. S1, S6B**). This template-based decoding task achieved 87±13% average accuracy for UHD-DOT but 74±17% average accuracy for HD-subset (**Figs. 6A-B**), a larger difference than for model-based decoding (*p* < 10^−4^, effect size 0.842). It is therefore likely that for model-based decoding, the breadth of the training set was a major limiting factor rather than image resolution, whereas for template-based decoding, image quality was the limiting factor. In other words, our decoder was “under-trained” for the model-based task, and we would expect the decoding performance to increase and the gap between UHD-DOT and HD-subset to widen if, in a follow-up study, a more-extensive, broader training set were to be used.

**Figure 6.**
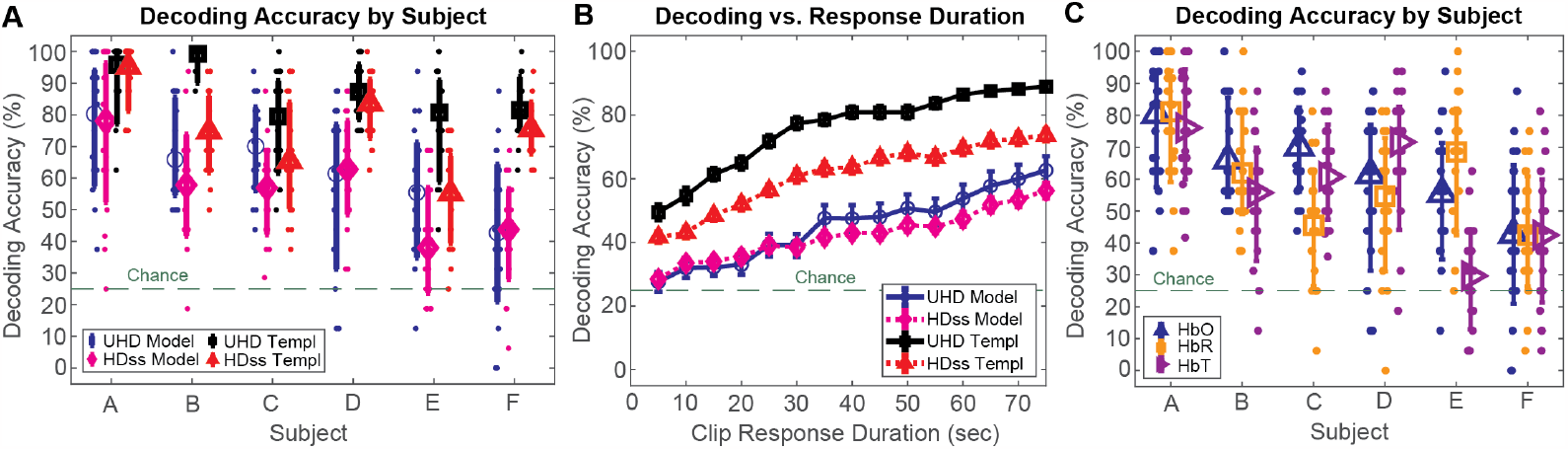
Decoding accuracy vs. imaging array, duration, and contrast. Imaging pipeline manipulations reveal impact of optode grid density and hemoglobin contrast type on decoding performance. (A) Model-based and template-based 4-clip decoding accuracy from UHD and HD-subset optode grids for each subject, session, and 6 train-test splits. Large markers and error bars denote mean ± lower and upper standard deviation over sessions and splits. (B) Decoding accuracy declines as expected as time window for decoding shortens. Error bars denote standard error over subjects, sessions, and splits. The denser grid gave a far-smaller improvement in model-based decoding than template-based decoding, suggesting that image quality was the limiting factor for template-based but not model-based decoding. (C) Model-based decoding accuracy for oxyhemoglobin (HbO), deoxyhemoglobin (HbR), and total hemoglobin (HbT) indicate statistical equivalence across these three contrasts, though HbO performed best in most subjects. Large markers and error bars denote mean ± lower and upper standard deviation over sessions and splits.

To examine whether the type of hemoglobin contrast would yield different decoding performances, we repeated our model training and our 4-clip decoding task with 2 test clips and 2 lures using deoxy-hemoglobin (HbR) and total hemoglobin (HbT) images. Oxy-hemoglobin (HbO) performed best, followed by HbR, and then HbT. Although these differences were statistically significant (*p*<0.01), their effect sizes were modest, with only the HbO-vs.-HbT effect size exceeding 0.2 (Cohen’s d 0.299, **Supplementary Table S2**).

## 3. DISCUSSION

Overall, the results demonstrate robust model-based decoding of naturalistic scenes from HD-DOT data. We evaluated decoding of naturalistic movie sequences that were outside the training set and found decoding accuracies that far-exceeded chance performance and that were robust to several manipulations. The decoding accuracy declines as the test-time window shortens, as expected, but remained well above chance for windows longer than 25 seconds (**Fig. 5B**). As the set of test clips was expanded, the decoding accuracy dropped, but the chance threshold dropped more quickly (**Fig. 5C**). Finally, we found that the training from one day could be used to decode data from another day, a critical capability needed for building decoders based on large training data sets (**Fig. 5B**). Most previous fNIRS decoding studies to date have accomplished coarse decoding among two or three very different but predetermined stimuli, conditions, or imagined actions.^9,26^ A few fNIRS decoders have accomplished 4-way decoding, but not beyond repeats of the decoder’s training set.^9,10^ These prior studies were neither model-based/parametric nor out-of-sample/extrapolative. The success of HD-DOT model-based decoding results herein represents a pivotal milestone for human optical neuroimaging and points toward HD-DOT’s promising potential for other advanced extrapolative decoding tasks.

In the last decade, using cortical hemodynamics, fMRI studies have progressed from identifying images with visual motion energy feature encoding models^2^ all the way to using semantic features and large language models to reconstruct natural language sentences.^5^ The results reported here represent the first step along a similar path for HD-DOT, leveraging a wavelet motion energy encoding model to decode naturalistic visual scenes, similar to the first important steps for fMRI.^1,2^

Model-based decoding allows integration of data from a diverse training set.^1–3,5,27^ Not only does that build a more-robust, more-capable decoder, but it also avoids repetitions that the subject may find boring or begin to ignore. The minimal decline in decoding performance when training the decoder with data from different imaging sessions (**Fig. 5B**) suggests that HD-DOT will be able to successfully integrate multi-session imaging data. For large training data sets, multi-session integration will be a requirement.

One straightforward extension from these results would be to greatly expand the amount of training data. While this study used a small training set for model-based decoding, with 3 minutes of unique video repeated 4-8 times, for a total of 12-24 minutes per subject, earlier fMRI studies have leveraged 120 minutes of unique video (40x more unique video than these results). Undertraining is reflected in the analysis of UHD versus HD subset (HDss) which generally shows improvement for UHD over HDss. However, the effect was greater with template-based decoding rather than with model-based decoding. For model-based decoding, rather than spatial resolution being a limit (UHD vs. HDss), it was the amount of training data. Decoding accuracy also declined but remained far above chance as the number of clips on the decoder’s menu increased, indicating the robustness and generalizability of this decoding approach. These results collectively suggest that our model decoder was undertrained by the reduced breadth/variety of the training set compared to previous fMRI studies.

The HD-DOT generalizability and scalability trends here echo those observed in fMRI visual decoding.^1,2^ DOT decoding is expected to scale in performance if the training set is expanded to the breadth of stimuli included in some of the pivotal fMRI decoding studies. If so, then these results may point to future success with the other fMRI decoding work, including visual semantics,^3^ auditory semantics, and finally sentence decoding.^5^ Scanning for two or more hours would be possible with HD-DOT. However, an experimental plan with data integration over multiple session would be more robust, and our multi-session results support that approach. Deep learning approaches^4,28^ will also improve and expand decoding, but were not evaluated here because our dataset did not provide the large, broad pool of training data they require.^29^

Another natural extension would be to move to a HD-DOT system that has full-head coverage. The field of view (FOV) of the UHD imaging system used here was restricted to occipital cortex, which mainly encodes visual feature information. While reasonable for a motion energy model, there are alternate models, for example semantic models of visual content, which build a decoding weight matrix that covers a much greater extent of the cortex.^3,5,30^ Our stimuli and FOV also did not leverage any auditory content. Auditory content, either as audio only,^5,30^ or integrated with visual content as with a normal movie,^3^ would expand the information available for training and decoding. Thus overall, an HD-DOT system with larger FOV would open up a greater variety of brain functions and enable more ambitious model construction.

Improving the response time of decoder is a reasonable goal for any brain computer interfaces.^6,7,31^ In this study, shorter test windows produced lower accuracy but remained far above chance, indicating that test trials need not be as long as 90 seconds to achieve meaningful decoding. Both extensions discussed above, larger training data and expanded FOV, have the potential to improve future decoding speed. Larger training data sets will provide a higher-fidelity feature-voxel map and increased signal to noise, while a larger FOV will bring more voxels into the decoding estimation calculation, providing higher accuracy from shorter time windows. We will still be left with the sluggishness of the hemodynamic response, which places a somewhat hard cap on decoding times at around 10 seconds. However, this apparent time constant limitation manifests in a variety of ways depending on the particulars of the decoding paradigm. If real-time control and <1-second precision is required, then hemoglobin responses to neural activity are likely not the right physiology to work with. However, if the goal is to decode a rich task, like language expression, and a 10-second delay is workable, there are new approaches that leverage large language models and work within the limitations of the hemodynamic response to provide reconstructed speech with word rates faster than one word per second.^5^

Our study represents a substantial advance in human optical brain decoding and suggests promising, expansive future directions. Larger training sets can improve the decoding accuracy of future studies, expand the range of stimuli that can be decoded, and broaden the range of decoding techniques that can be employed. A larger field of view will open more decoding possibilities, such as harnessing semantic and/or auditory information for category and sentence decoding. Furthermore, these model-based decoding techniques are expected to work with emerging wearable HD-DOT systems^19–21^ to bring powerful, expansive decoding capabilities into naturalistic, everyday settings and longer-term clinical uses such as brain-computer interface.

## METHODS

### M1. Visual Stimuli, Protocol, and Subjects

To evaluate brain response repeatability to movie clips and evaluate decoding among at least 4 clips, we showed 12 subjects four 90-second clips of interest in random order 4-10 times per imaging session in 4 sessions. We selected clips from a successful fMRI encoding and decoding study,2 and clips contained no audio. All movie clips and movie images were accessed and presented from CRCNS.org32 by Washington University in St. Louis-affiliated authors. Subjects were healthy adults, with 5 females, 7 males, and age range 24–31 years. We restricted most analyses to the 6 subjects with adequately repeatable brain responses to each clip to avoid confounding effects from some subjects insufficiently attending to the stimuli or failing to fixate on the center of the screen (**Supplementary Figs. S1-S3**).

### M2. Imaging System, Data Preprocessing, and Raw Data Quality Control

To image hemodynamic signals from occipital brain areas sensitive to visual stimuli, we placed an ultra-high-density DOT system with 126 sources and 126 detectors covering a 12cm x 5.5cm area on the back of the head,^22^ and we employed well-established signal processing and image reconstruction pipelines for DOT.^13,15,33,34^ Briefly, we converted measured light level time traces in each measurement channel (i.e., each source-detector pair and light wavelength) into log-ratio time traces representing the fractional change in light level about its mean in that channel. Channels whose log-ratio time traces’ standard deviation over time exceeded 0.075 were labeled as excessively noisy and excluded from further analyses. We computed the signal-to-noise ratio of the cardiac pulse oscillations in each time trace as a measure of data quality, and we then applied a 0.02-0.2 Hz band-pass filter and superficial signal regression^35^ to the time traces to remove systemic signal components unrelated to the brain’s stimulus-evoked responses, such as cardiac pulse, respiration, scalp dynamics, and brain-wide blood oxygen and volume changes. Next, we reconstructed images of the absorption coefficient fluctuation from baseline at each time point by applying a spatially-variant Tikhonov (L2)-regularized pseudoinverse operator to the preprocessed channels at each time point. The regularized pseudoinverse was computed from a sensitivity matrix that expresses the channels’ preprocessed measurements at each time point as a weighted sum of absorption coefficient fluctuations at each location (voxel) in the head. That sensitivity matrix was computed from finite-element simulations of light propagation through the head of a single subject, a segmented 5-layer model of scalp, skull, cerebrospinal fluid, gray matter, and white matter tissue obtained from anatomical T1-weighted and T2-weighted MRI head images acquired in a previous study.^22^ Next, the absorption coefficient fluctuation images at each light wavelength were spectroscopically converted into concentration fluctuations in oxyhemoglobin and deoxyhemoglobin. Finally, to further isolate brain-specific signals, voxels in scalp and skull were excluded from encoding model fitting and clip identity decoding analyses. We employed the NeuroDOT^33,34^ and NIRFAST^36^ software packages for these processing steps.

### M3. Encoding Model Features

To decompose any clip into a common set of features to which occipital cortex areas respond strongly, we computed the wavelet motion energy in a sliding 4-second time window at multiple locations, scales, spatial frequencies, and temporal frequencies in each clip (**Supplementary Fig. S4**). These same features enabled accurate predictive encoding models and decoding in a previous fMRI study,^2^ and we adapted the feature decomposition pipeline^37^ to include a canonical hemodynamic response function^38^ and a band-pass filter matching the DOT preprocessing pipeline.^13^

### M4. Encoding Model Fitting and Prediction Performance

To estimate the encoding model weights, we performed ridge regression on the DOT images and stimulus motion energy features for the training clips (**Fig. 2**). During fitting, we employed 4-fold cross-validation on the training data, swept the regression regularization parameter, and selected the value that yielded highest average prediction accuracy on the validation portion of the training data. We then applied that optimal regularization parameter to estimate the model weights from the whole training set. The test set was completely held out from this fitting procedure.

To evaluate how well the model predicted brain responses, we computed the correlations between the predicted responses to each non-training clip and the measured responses to the same clips (“matched-clip” case) or to a gray fixation cross (“resting state”). We computed a Cohen’s d statistic^39^ as a contrast-to-noise ratio (CNR) measure of the separation between the matched-clip correlations and clip-to-rest correlations, which quantifies the strength and reliability of the encoding model’s predictions above a “null”/mismatched case (**Supplementary Fig. S5**).

### M5. Model-Based Decoding

To identify clips outside the training set, we used the encoding model to predict the response to each clip in a “menu” including the test clips, and for each trial, our decoder guessed that the subject was viewing the clip whose predicted response was most highly correlated with the measured response (**Fig. 4C, Supplementary Fig. S6**). Decoding accuracy was quantified by the percentage of trials in which the decoder identified the correct clip.

To evaluate the encoding model’s ability to support cross-session decoding, we trained the encoding model on one session and used it to predict responses for decoding in other sessions within each subject. To evaluate the impact of using shorter durations of data for encoding and decoding, we repeated our fitting and decoding procedures on progressively shorter time windows of each clip and response (**Figs. 5B, 6B**). To evaluate generalizability to many-clip decoding, we gradually increased the number of lure clips whose predicted responses were included on the decoder’s “menu” (**Fig. 5C**).

To assess the impact of imaging grid density, we repeated several decoding analyses on images reconstructed from a sparser subset of the DOT measurement channels corresponding to a more-standard high-density DOT system (**Figs. 6A, 6B**). We also repeated several decoding analyses with deoxy- and total-hemoglobin images to evaluate whether the choice of hemoglobin contrast would affect decoding accuracy (**Fig. 6C**).

### M6. Template-Based Decoding

To perform a more-basic, template-based decoding task similar to previous HD-DOT decoding^18,22^ as a validation and comparison, we treated the data from the first and second halves of each imaging session as training and testing sets, then block-averaged the responses to each clip to serve as “templates,” then had a decoder guess that the clip being viewed was the one whose template response was most highly correlated with the actual response in each test trial. We repeated many of the manipulations described elsewhere in this paper to evaluate template-based decoding performance for between-session training and testing, decreasing time window duration, increasing number of clips to decode, and reducing optode grid density (**Fig. 6A-B; Supplementary Figs. S6B, S7**).

### M7. Statistical Information

To test statistical significance of decoding accuracies against chance level, we performed a binomial test. To compare matched-clip vs. rest correlations, within-session vs. between-session decoding performance, and template-based vs. model-based decoding performance, we employed independent two-sample bootstrap tests for statistical significance.^40^ For all other statistical comparisons, we employed matched/paired two-sample permutation testing for significance.^41^ Each bootstrap and permutation test employed 10^4^ iterations/permutations. We employed Cohen’s d statistic^39^ to measure effect size in all statistical comparisons. More-detailed statistical information is provided in the Supplementary Information (**Section S8**).

### M8. Ethical Approval for Human Experiments

Informed consent was obtained from all participants. Research was approved by the Washington University Institutional Review Board (IRB) under IRB protocol #201707092 and was performed in accordance with this protocol.

## AUTHOR CONTRIBUTIONS

Z.E.M., K.T., E.J.R., A.T.E., M.A.A., M.A.C., E.M.M., S.N.N., A.Y., J.W.T., and J.P.C. conceived and designed the experiments. Z.E.M., K.T., A.M.S., M.L.S., S.M.R., and J.W.T. performed the experiments. Z.E.M., K.T., A.M.S., M.L.S., A.T.E., M.A.A., M.A.C., E.M.M., S.N.N., A.Y., J.W.T., and J.P.C. analyzed the data. Z.E.M., K.T., J.W.T., and J.P.C. contributed materials/analysis tools. Z.E.M., K.T., J.W.T., and J.P.C. wrote the paper.

## ACKNOWLEDGMENTS

The authors thank Patrick Mineault and Michael A. Choma for helpful conversations about this work. We also gratefully acknowledge funding from the following sources: a Meta Sponsored Academic Research Agreement (J.P.C.), National Institutes of Health grants: U01EB027005 (J.P.C.), R01NS090874 (J.P.C.), R01EB034919 (J.P.C.), K01MH103594 (A.T.E.), R01MH122751 (A.T.E.), and F31NS110261 (Z.E.M.) and Washington University’s Cognitive, Computational, and Systems Neuroscience Fellowships (Z.E.M., K.T.).

## DATA AVAILABILITY AND CODE AVAILABILITY

Data collected and analyzed in this study are available from the corresponding authors upon reasonable request. Software code employed in this study is available from the corresponding authors upon reasonable request.

