## Supplementary Information for "Identifying Naturalistic Movies from Human Brain Activity with High-Density Diffuse Optical Tomography"

### S1. Train-test splits for repeatability analysis and decoding tasks

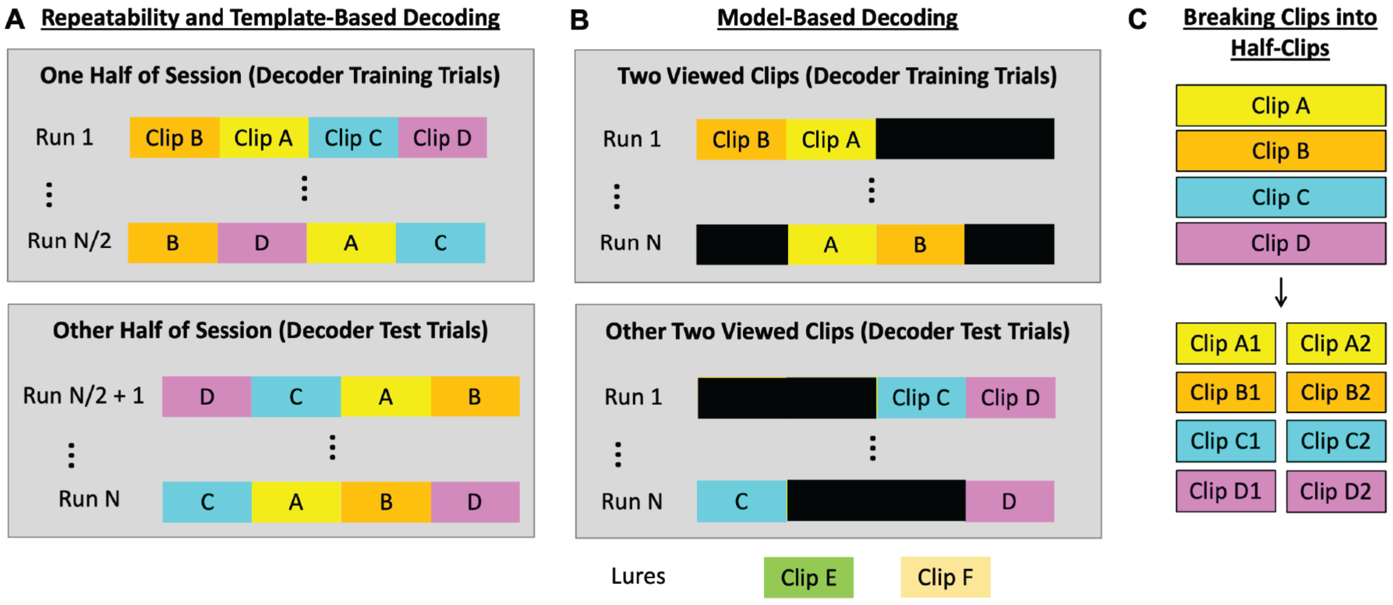


**Supplementary Figure S1:** Example train-test splits of clip responses from each imaging session for repeatability analysis, template-based decoding, and model-based decoding. (A) To evaluate clip response repeatability and perform template-based decoding, the responses to all clips from half the runs in an imaging session were treated as training trials, and the remaining clip responses were treated as test trials. (B) To train the encoding model and perform model-based decoding, two clips and their responses from all runs in an imaging session were treated as training data, and the other two clips and their responses were treated as test data. For some analyses, the responses to never-viewed "lure" clips were predicted and included in the decoder's menu of options to make decoding more challenging. (C) For some analyses, including Figure 4, each clip was treated as two successive "half-clips", and each trial was similarly treated as two successive half-length trials. For the train-test split in Figure 4, the four test half-clips were Clips A1, A2, C1, and C2.

### S2. Clip response repeatability contrast-to-noise ratio (CNR)

To quantify the repeatability of the brain's response to each clip in each subject and session, we used a contrast-to-noise ratio (CNR) metric measuring the statistical separation between a "matched-clip" and a "rest" (null) distribution of transformed correlations. To calculate this CNR, we block-averaged the response to each clip from one half of the runs within a session. We then calculated the spatiotemporal correlation between these mean responses and the single-trial responses to the same clips in the other half of the session to form the "matched-clip" distribution. We also calculated the spatiotemporal correlation between these mean responses and resting-state activity periods of equal duration to form a "rest" (null) distribution. We also applied Fisher's r-to-z transformation[1-3] to each correlation value to make their distributions more Gaussian-like. We then calculated Cohen's d statistic[4] between these matched-clip and rest distributions as the measure of CNR (**Supplementary**

**Fig. S2**). Thus the repeatability CNR value is the number of standard deviations by which the transformed matched-clip correlations exceed clip-to-rest correlations on average.

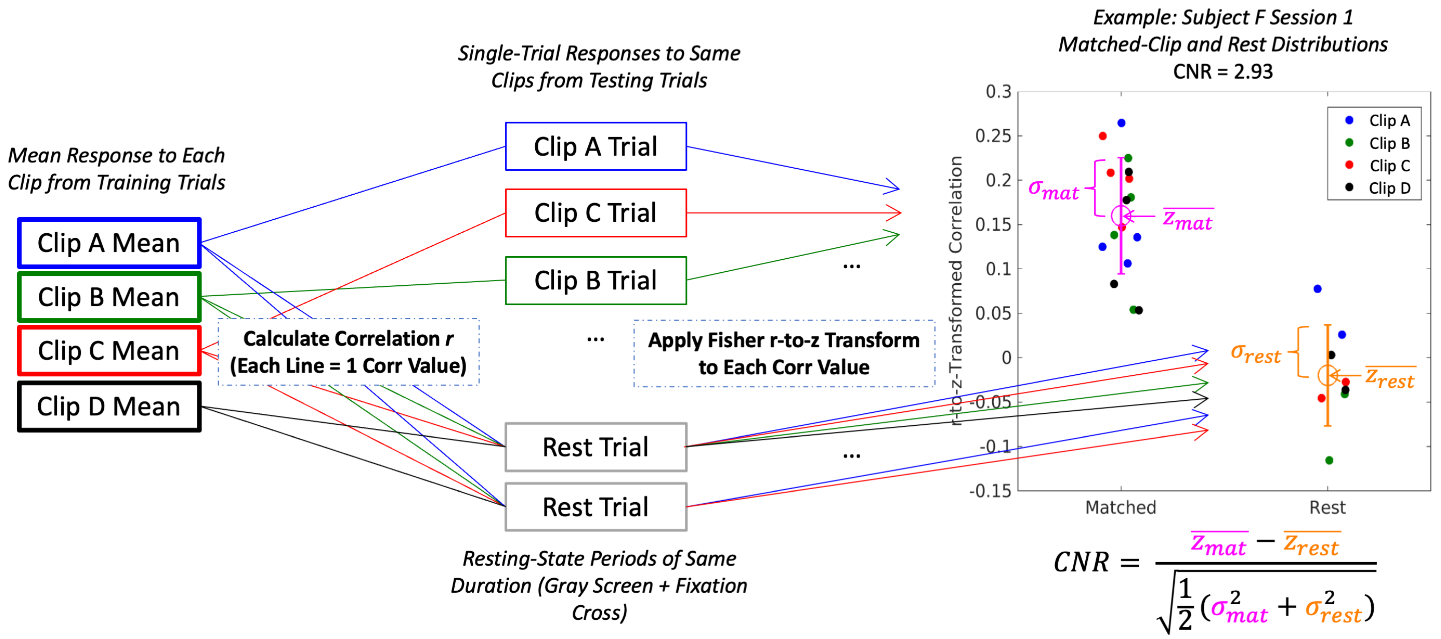


**Supplementary Figure S2:** Quantifying clip response repeatability. We block-averaged the brain responses to each clip in one half of an imaging session to form mean responses. We then calculated the spatiotemporal correlations between each mean response and the single-trial responses from the same clip in the other half of the data and Fisher's-r-to-z-transformed these correlations to form a matched-clip distribution. We also calculated the spatiotemporal correlations between each mean response and the brain activity in resting-state periods of equal duration and applied the same Fisher's-r-to-z transformation to form a "rest"/null distribution of correlations against which to compare the matched-clip distribution. We then quantified repeatability CNR with Cohen's d statistic, which measures the separation of these distributions' means relative to their standard deviations. An example of the distributions and CNR calculation is shown on the graph for one training-test split of the trials for one session in one subject.

### S3. Clip response repeatability as a data quality control metric

Successful, reliable decoding requires that a subject's neural responses to a given movie clip be repeatable. Extraneous behavioral factors such as low attention to the movie clips or failure to fixate consistently on the central dot could reduce the repeatability of a subject's evoked brain responses to each movie clip and reduce decoding performance. Therefore, we restricted most analyses to the 6 subjects whose average clip repeatability CNR was no less than 1.5 over sessions and training-test splits. As expected, there were strong positive correlations between clip response repeatability CNR vs. model-based and template-based decoding performance (0.834 for model-based and 0.884 for template-based decoding).

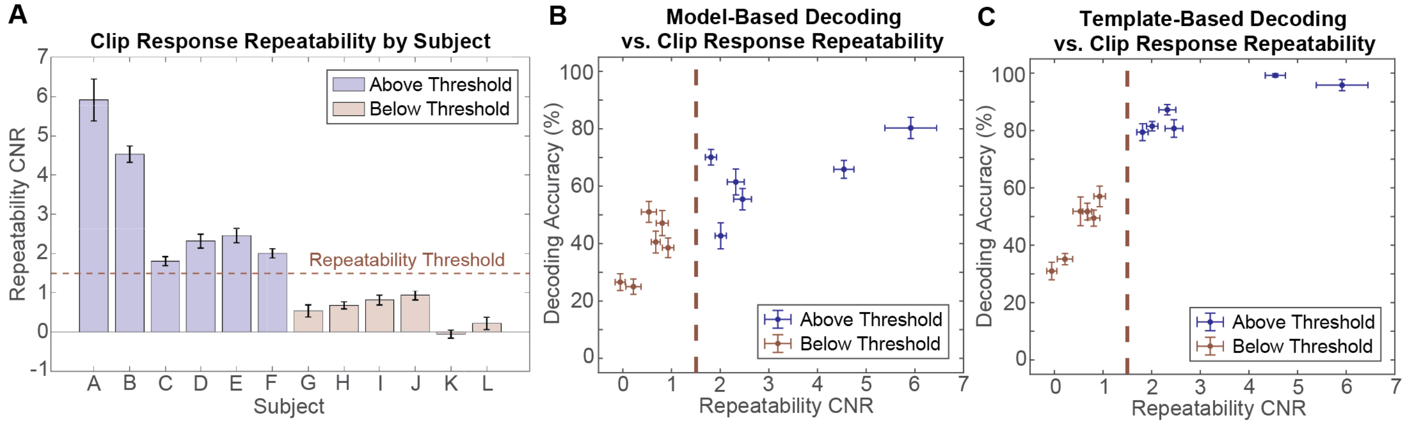


**Supplementary Figure S3:** Clip response repeatability as a data quality control metric. (A) Average clip response repeatability CNR over sessions and training-test splits within each subject. Error bars denote standard error of the mean. A repeatability CNR threshold of 1.5 (dashed horizontal red line) was imposed to exclude subjects whose brain responses to repeated presentations of each clip were less than 1.5 standard deviations more correlated than to resting-state periods, in which no movie clip was shown. (B-C) Model-based and template-based decoding accuracy were both strongly correlated with clip response repeatability. Error bars denote standard error of the mean. The dashed vertical red line shows the repeatability CNR threshold (1.5).

### S4. Wavelet motion energy calculation

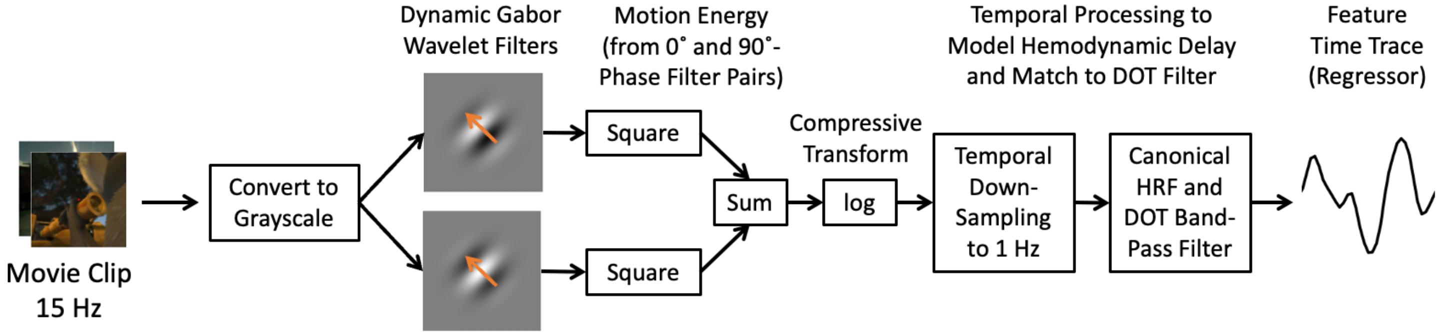


**Supplementary Figure S4:** Pipeline for decomposing any movie clip into wavelet motion energy feature time traces. Each wavelet motion energy feature quantifies the activity at a given spatial frequency, speed, direction, and location in the movie clip. These energy traces were filtered to account for the hemodynamic response delay[5] and match the frequency content of the HD-DOT imaging data. This pipeline was closely based on the successful fMRI decoding study from which the stimuli were taken[6] and employed the related "motion_energy_matlab" software package.[7]

### S5. Quantifying encoding model prediction accuracy

To quantify the accuracy with which the encoding model predicted the brain's responses to the clips, we calculated the spatiotemporal correlation between the measured and predicted brain responses in each test trial. For comparison, we also calculated the spatiotemporal correlation between the predicted brain responses and measured brain activity during resting-state periods of equal duration. We then Fisher-r-to-z-transformed these correlation distributions and compared them using Cohen's d, much like how we calculated clip response repeatability CNR.

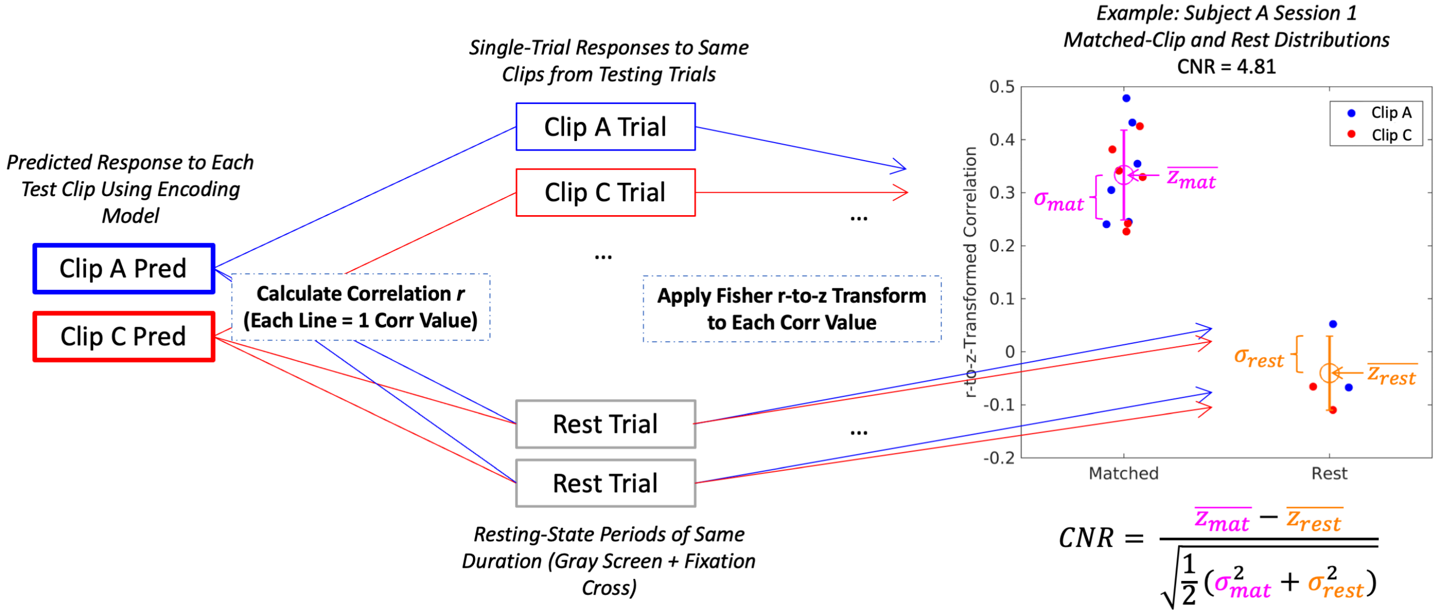


$$CNR = \frac{\overline{z_{mat}} - \overline{z_{rest}}}{\sqrt{\frac{1}{2}(\sigma_{mat}^2 + \sigma_{rest}^2)}}$$

**Supplementary Figure S5:** Quantifying encoding model prediction accuracy. We used the encoding model to predict the brain's responses to each test clip. We then calculated the spatiotemporal correlations between each predicted response and the measured single-trial responses from the same clips. We Fisher's-r-to-z-transformed these correlations to form a matched-clip distribution. We also calculated the spatiotemporal correlations between each predicted response and the brain activity in resting-state periods of equal duration and applied the same Fisher's-r-to-z transformation to form a "rest"/null distribution of correlations against which to compare the matched-clip distribution. We then quantified prediction accuracy "CNR" with Cohen's d statistic. An example of the distributions and prediction accuracy CNR calculation is shown on the graph for one training-test split of the clips for one session in one subject.

### S6. Maximum-correlation decoding procedure

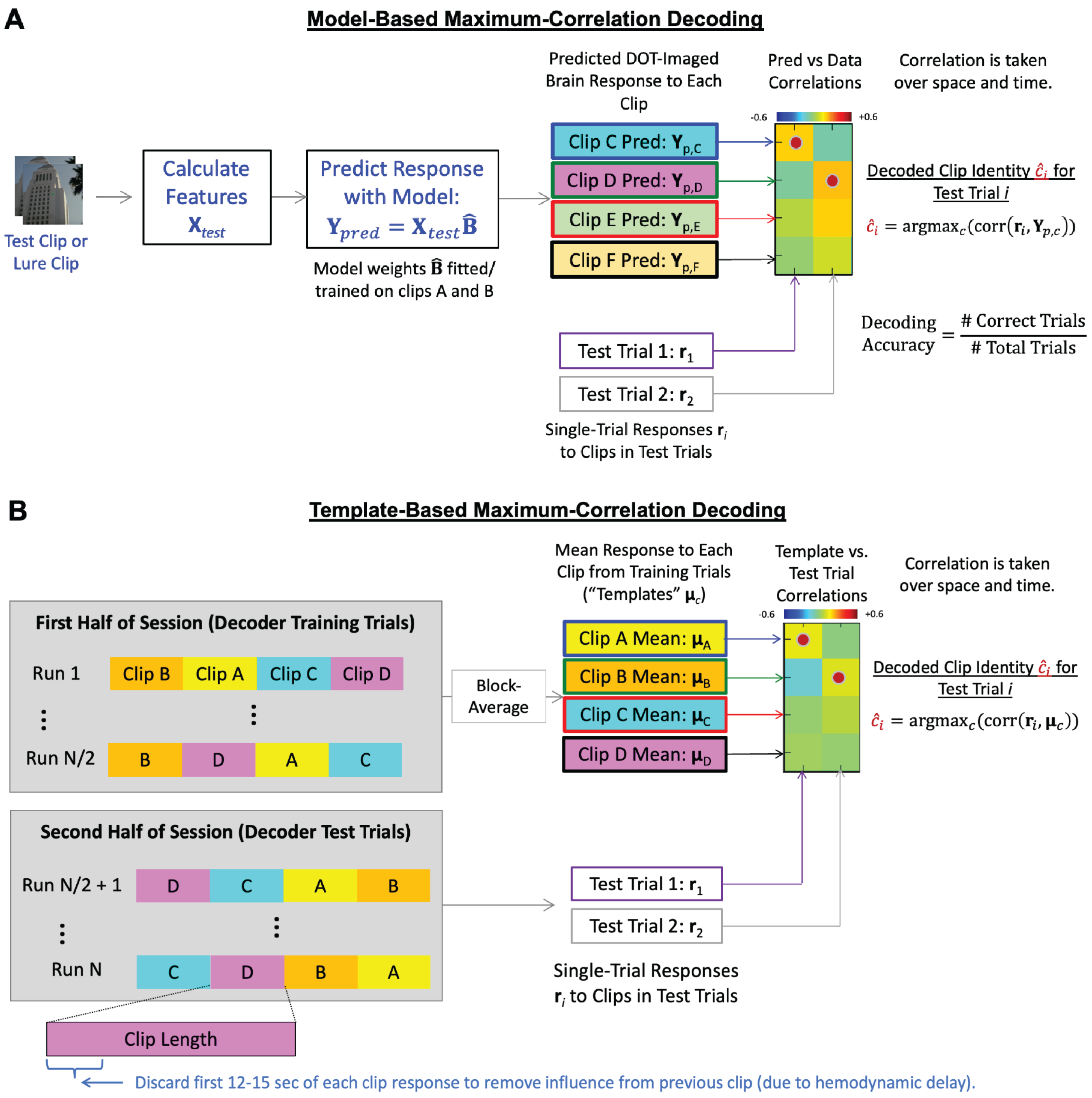


**Supplementary Figure S6:** (A) To perform model-based decoding, we calculate the brain's predicted response to each test and lure clip, and then we calculate the correlation over space and time between each predicted response and the actual, measured response in each test trial, forming a predicted responses x test trials correlation matrix. In each trial, the decoder guesses that the clip being viewed was the clip whose predicted response was most correlated with the measured response. (B) To perform template-based decoding, we employ a similar procedure, but the correlations are calculated between each test trial's measured response and the block-averaged response ("template") for each clip from the training set instead of the model-predicted response to each clip. In both model-based and template-based decoding, the first 12-15 seconds of each trial is discarded to remove any spill-over influence from the previous clip due to the hemodynamic response delay.[5] 12 seconds are discarded when decoding among half-clips, and 15 seconds are discarded when decoding among full clips.

### S7. Template-based decoding between sessions and with more clips

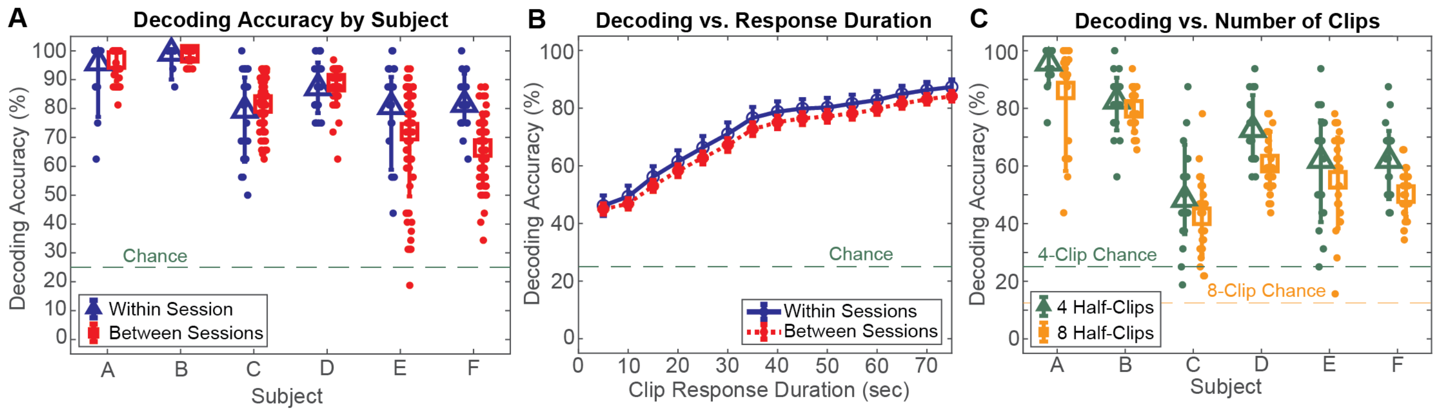


**Supplementary Figure S7:** Paradigm manipulations indicate decoder robustness and generalizability across sessions, data duration, and number of clips for template-based decoding. (A) Within-session and between-session decoding accuracy over sessions (for within-session) or session pairs (for between-session) and 6 train-test splits. Large markers and error bars denote mean ± lower and upper standard deviation. On average over sessions, subjects, and train-test splits, within-session and between-session template-based decoding accuracies were 87±13% and 84±16%, respectively, both far above chance ($p$<10$^{-5}$, effect sizes > 3.5). (B) Decoding accuracy declines as expected but remains above chance as the time window used for decoding shortens. Markers and error bars denote mean ± standard error over subjects, sessions, and 6 train-test splits. (C) Decoding accuracy declines but remains above chance, far above chance for some subjects, when the number of clips is doubled. Here, the 4 main clips and their responses were split in half for the 4-half-clip and 8-half-clip decoding, so the clip response duration used for decoding was identical for the 4-half-clip and 8-half-clip cases. Error bars denote lower and upper standard deviation over sessions and train-test splits. On average over sessions, subjects, and train-test splits, 4-half-clip and 8-half-clip template-based decoding accuracies were 69±20% and 62±20%, respectively, both far above chance ($p$<10$^{-5}$, effect sizes > 2.0).

### S8. Detailed statistical information

| *Decoding Condition* | *Related Figure(s)* | *Pooled Decoding Accuracy $a_P$ ± Std. Dev. s (%)* | *Chance-Level Decoding Accuracy $a_C$ (%)* | *95% Confidence Interval* | *Degrees of Freedom* | *Binomial Test p-Value* | *Effect Size (Cohen's d)* |
|---|---|---|---|---|---|---|---|
| Model-Based, 4 Test Half-Clips, Within Session, UHD Grid, HbO | 4D (orange triangles) | 53.9 ± 16.4 | 25.0 | [52.4, 55.3] | 4392 | $<10^{-5}$ | 1.76 |
| Model-Based, 2 Test Clips + 2 Lures, Within Session, UHD Grid, HbO | 4D (blue squares) | 62.0 ± 21.6 | 25.0 | [60.0, 64.1] | 2196 | $<10^{-5}$ | 1.71 |
| Model-Based, 2 Test Clips + 2 Lures, Between Sessions, UHD Grid, HbO | 5A (red squares) | 60.3 ± 22.6 | 25.0 | [59.2, 61.5] | 6588 | $<10^{-5}$ | 1.55 |
| Model-Based, 2 Test Clips + 2 Lures, Within Session, HDss Grid, HbO | 6A (magenta diamonds) | 55.5 ± 21.1 | 25.0 | [53.4, 57.6] | 2196 | $<10^{-5}$ | 1.45 |
| Model-Based, 2 Test Clips + 2 Lures, Within Session, UHD Grid, HbR | 6C (orange squares) | 58.3 ± 22.3 | 25.0 | [56.3, 60.4] | 2196 | $<10^{-5}$ | 1.49 |
| Model-Based, 2 Test Clips + 2 Lures, Within Session, UHD Grid, HbT | 6C (purple triangles) | 55.3 ± 23.4 | 25.0 | [53.2, 57.4] | 2196 | $<10^{-5}$ | 1.30 |
| Template-Based, 4 Clips, Within Session, UHD Grid, HbO | 6A (black squares) | 87.1 ± 13.0 | 25.0 | [85.7, 88.5] | 2208 | $<10^{-5}$ | 4.79 |
| Template-Based, 4 Clips, Within Session, HDss Grid, HbO | 6A (red triangles) | 74.2 ± 17.4 | 25.0 | [72.4, 76.1] | 2208 | $<10^{-5}$ | 2.84 |
| Template-Based, 4 Clips, Between Sessions, UHD Grid, HbO | S7A (red squares) | 83.6 ± 15.6 | 25.0 | [83.0, 84.3] | 13176 | $<10^{-5}$ | 3.76 |
| Template-Based, 4 Half-Clips, Within Session, UHD Grid, HbO | S7C (green triangles) | 69.5 ± 19.7 | 25.0 | [67.6, 71.4] | 2208 | $<10^{-5}$ | 2.26 |
| Template-Based, 8 Half-Clips, Within Session, UHD Grid, HbO | S7C (orange squares) | 61.8 ± 20.0 | 12.5 | [60.4, 63.3] | 4416 | $<10^{-5}$ | 2.46 |

**Table S1:** Detailed statistical information for comparing decoding accuracy against chance level in different decoding conditions. Pooled decoding accuracy $a_P$ is the number of correct test trials divided by the total number of test trials across all subjects, which is equal to the degrees of freedom for the corresponding binomial distribution. Binomial significance tests and confidence intervals were employed here because decoding accuracy is a proportion of successful trials, and the total number of trials was large enough to approximate the binomial distribution as Gaussian.[8] The standard deviation *s* in decoding accuracy was computed over 6 train-test splits and the 4 sessions in the 6 subjects with adequate clip response repeatability for inclusion in final analyses. Cohen's d in this case was calculated as $(a_P - a_C)/s$.

| *Decoding Conditions Being Compared* | *Related Figure(s)* | *Difference in Pooled Decoding Accuracy (%)* | *95% Confidence Interval* | *Degrees of Freedom* | *Significance Test p-Value* | *Effect Size (Cohen's d)* |
|---|---|---|---|---|---|---|
| Model-Based, 2 Test Clips + 2 Lures, UHD Grid, HbO, Within Session vs. Between Sessions | 5A (blue triangles - red squares) | 1.68 | [-0.66, 4.03] | 8784 | 0.0864 | 0.076 |
| Model-Based, 2 Test Clips + 2 Lures, Within Session, HbO, UHD Grid vs. HDss Grid | 6A (blue circles - magenta diamonds) | 6.51 | [3.61, 9.42] | 2196 | $<10^{-4}$ | 0.305 |
| Template-Based, 4 Clips, Within Session, HbO, UHD Grid vs. HDss Grid | 6A (black squares - red triangles) | 12.9 | [10.6, 15.2] | 2196 | $<10^{-4}$ | 0.842 |
| Template-Based 4 Clips vs. Model-Based 2 Test Clips + 2 Lures, HbO, UHD Grid | 6A (black squares - blue circles) | 25.1 | [22.7, 27.6] | 4404 | $<10^{-4}$ | 1.41 |
| Template-Based 4 Clips vs. Model-Based 2 Test Clips + 2 Lures, HbO, HDss Grid | 6A (red triangles - magenta diamonds) | 18.7 | [16.0, 21.5] | 4404 | $<10^{-4}$ | 0.970 |
| Model-Based, 2 Test Clips + 2 Lures, Within Session, UHD Grid, HbO vs. HbR | 6C (blue triangles - orange squares) | 3.69 | [0.79, 6.58] | 2196 | $10^{-4}$ | 0.168 |
| Model-Based, 2 Test Clips + 2 Lures, Within Session, UHD Grid, HbO vs. HbT | 6C (blue upward triangles - purple rightward triangles) | 6.74 | [3.83, 9.65] | 2196 | $<10^{-4}$ | 0.299 |
| Model-Based, 2 Test Clips + 2 Lures, Within Session, UHD Grid, HbR vs. HbT | 6C (orange squares - purple triangles) | 3.05 | [0.12, 5.98] | 2196 | 0.0086 | 0.135 |

**Table S2:** Detailed statistical information for comparing decoding accuracy pairwise between decoding conditions. We evaluated statistical significance with matched-pairs permutation tests except for comparing within-session vs. between-session decoding and template-based vs. model-based decoding, for which we employed 2-independent-samples bootstrap tests. Permutation and bootstrap tests each employed $10^4$ permutations/iterations. Gaussian approximations to the binomial confidence intervals are reported due to the large number of trials. All these values were computed over 6 train-test splits and the 4 sessions in the 6 subjects with adequate clip response repeatability for inclusion in final analyses.

| *Distributions Being Compared* | *Related Figure(s)* | *Pooled Difference* | *95% Confidence Interval* | *Degrees of Freedom* | *Significance Test p-Value* | *Effect Size (Cohen's d)* |
|---|---|---|---|---|---|---|
| Model Prediction-to-Measured Response Correlations vs. Model Prediction-to-Rest Response Correlations ("Matched-Clip" vs. "Rest" Distributions) | 3E-G, S5 | 0.100 | [0.098, 0.103] | 13390 | $<10^{-4}$ | 1.31 |

**Table S3:** Detailed statistical information for evaluating encoding model prediction accuracy. The "matched-clip" vs. "rest" distributions of transformed correlation values described in Supplementary Information Section S5 were pooled for 6 train-test splits and the 4 sessions in the 6 subjects with adequate clip response repeatability for inclusion in final analyses. These statistics were computed on these two pooled distributions. The confidence interval was computed from Student's *t* distribution, and significance was determined by a 2-independent-samples bootstrap test with $10^4$ permutations/iterations.

Supplementary Information References


1 Fisher, R. A. Frequency Distribution of the Values of the Correlation Coefficient in Samples from an Indefinitely Large Population. *Biometrika* **10**, 507-521 (1915). https://doi.org:10.2307/2331838

2 Fisher, R. A. On the 'probable error' of a coefficient of correlation deduced from a small sample. *Metron* **1**, 1-32 (1921).

3 Hotelling, H. New Light on the Correlation Coefficient and its Transforms. *Journal of the Royal Statistical Society. Series B (Methodological)* **15**, 193-232 (1953).

4 Cohen, J. *Statistical Power Analysis for the Behavioral Sciences*. 2nd edn, (Lawrence Erlbaum Associates, 1988).

5 Hassanpour, M. S. *et al.* Statistical analysis of high density diffuse optical tomography. *Neuroimage* **85 Pt 1**, 104-116 (2014). https://doi.org:10.1016/j.neuroimage.2013.05.105

6 Nishimoto, S. *et al.* Reconstructing visual experiences from brain activity evoked by natural movies. *Curr Biol* **21**, 1641-1646 (2011). https://doi.org:10.1016/j.cub.2011.08.031

7 Nishimoto, S. (https://github.com/gallantlab/motion_energy_matlab, GitHub, 2011).

8 Tamhane, A. C. & Dunlop, D. D. *Statistics and Data Analysis from Elementary to Intermediate*. (Prentice Hall, 2000).
